# The development of efficient communication in the human connectome

**DOI:** 10.1101/2022.03.14.484297

**Authors:** D. van Blooijs, M.A. van den Boom, J.F. van der Aar, G.J.M. Huiskamp, G. Castegnaro, M. Demuru, W.J.E.M. Zweiphenning, P. van Eijsden, K. J. Miller, F.S.S. Leijten, D. Hermes

## Abstract

The structure of the human connectome develops from childhood throughout adolescence to middle age, but how this affects the speed of neuronal signaling is not well described. To understand the maturation process of transmission speed across the connectome, we stimulated electrocorticographic (ECoG) electrode pairs in 75 human subjects of 4 to 51 years old. We then measured the latency of cortico-cortical evoked responses in other electrodes. Each year, responses got faster, until transmission speed reached a minimum latency at the age of 29-44 years old. Responses during middle age were about two times faster compared to childhood. These increases in speed were observed in both long- and short-range connections between and within frontal, central, parietal, and temporal areas. These findings indicate that long-range and short-range communication between brain regions increase in efficiency well throughout adolescence.

## Introduction

The development of rapid communication between human brain regions is essential for cognitive function. The speed of neuronal transmission is fundamental to many computational human brain models (*1*), as it influences whether electrical signals arrive at the same or at different times. This directly affects the timescales across which information is integrated (*2*) and the amount of time it takes to reach a decision (*3, 4*). However, little is known about the maturation process of transmission speed in the human brain, partially because the axonal diameter in the adult human brain is relatively large compared to most other mammalian species (*5*).

Rapid neuronal communication in the human brain may be largely established during early childhood. EEG studies, measuring the latency of visual and auditory evoked potentials as an approximation of transmission speed, find major changes during infancy (*6*). Some EEG studies have detected small changes during adolescence (*7, 8*). After adolescence there may be little further maturation, as also indicated by one study of cortico-cortical evoked potentials (CCEPs), which reported a decrease in conduction delay of only one millisecond from before to after the age of 15 (*9*).

Anatomical studies, however, indicate that the human connectome may follow a much longer developmental trajectory: postmortem studies have shown that myelination starts in the late prenatal period and continues into late adolescence (*10*). MRI analyses have demonstrated that white matter properties change across the life-span (*11*), often reaching a plateau around 30 years of age (*12, 13*), well beyond adolescence. In this study, we investigate whether this known structural development translates to changes in the timescales of neuronal communication.

## Results

To characterize the timescales of transmission speed in the human connectome, we measured CCEPs during human intracranial electrocorticography (ECoG) recordings. CCEPs were measured for clinical purposes in a large group of 75 subjects (age 4-51 years old) who had ECoG electrodes implanted subdurally for epilepsy monitoring (Figure 1A). Adjacent electrode pairs were stimulated with ten single monophasic electrical pulses of 1 ms width at 0.2 Hz. Amplitudes varied from 4-8 mA (which did not systematically influence the results, see Fig S1). Responses to stimulations in other electrodes were averaged across the ten pulses, excluding channels and epochs with artifacts. CCEPs often show an early surface negative deflection (N1) within 100 ms after stimulating another electrode pair (Figure 1B). Frontal and parietal regions are connected through the superior longitudinal fasciculus and Figure 1B shows an example where the N1 response measured in frontal areas upon parietal stimulation peaks around 45 ms in a 4-year-old. In a 38-year-old, however, the N1 response peaks about 1.5 times faster, around 30 ms.

**Fig. 1.**
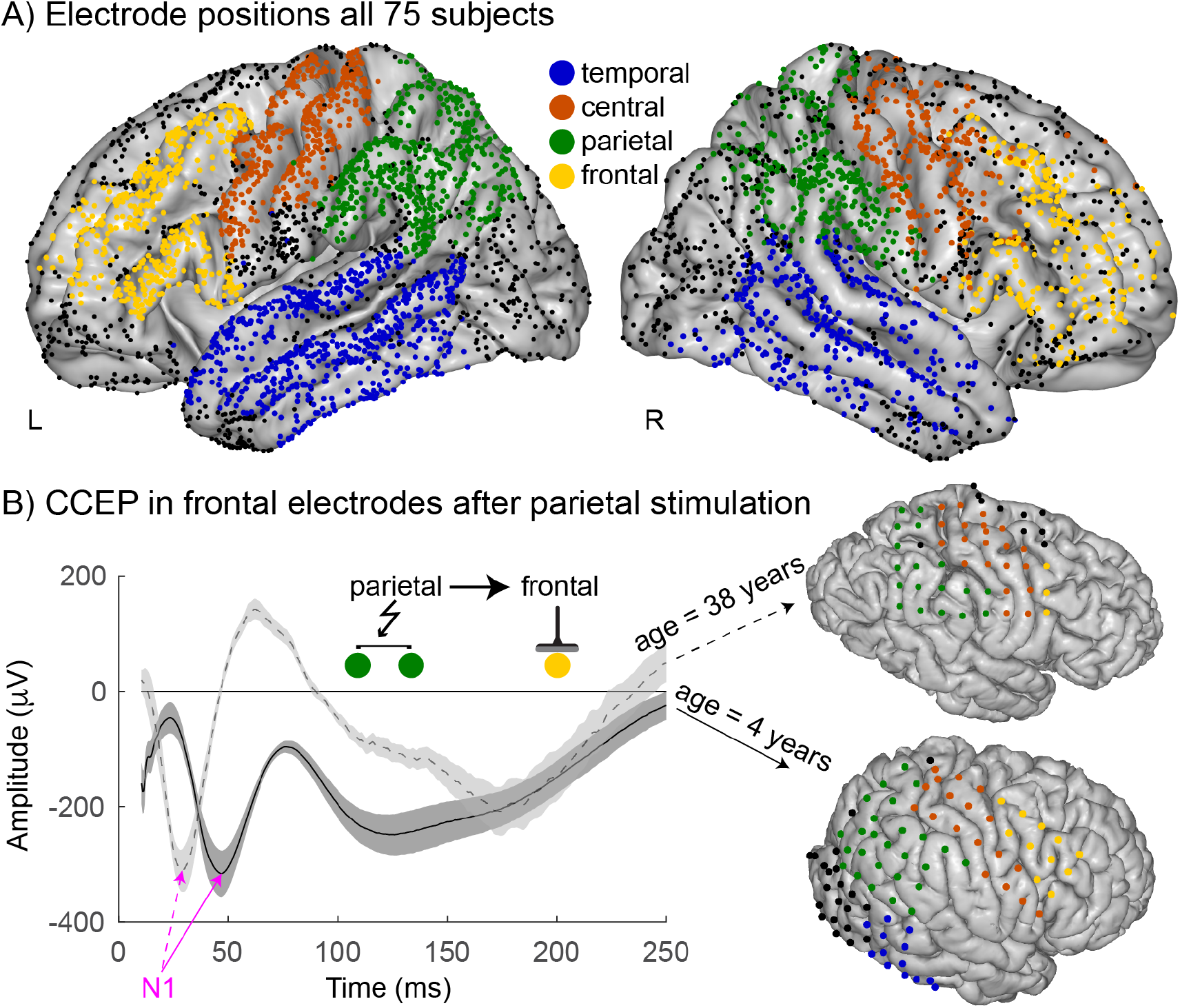
Electrode positions and evoked potentials. (**A)** Standardized brain with electrode positions from all 75 subjects. Temporal (blue), central (red), parietal (green) and frontal (yellow) electrodes labeled using Freesurfer (18) and included in analyses. (**B)** Left: CCEPs (+/- s.e.) from a 4-year-old (solid) and a 38-year-old (dashed) averaged across responding frontal electrodes after parietal stimulation showing an N1 peak (magenta). Right: individual brain rendering with electrodes.

This rapid surface negative N1 potential has been related to direct cortico-cortical connections (*14*) and is thought to be generated by synchronized, excitatory synaptic activation of the distal layer apical dendrites of the pyramidal cells (*15*). While this feature selection likely ignores many other aspects of the evoked potential that provide a richer characterization of cortico-cortical communication (*16*), the N1 response provides insight into transmission speed across the human connectome (*9, 17*).

To quantify changes in transmission speed across the group, we calculate the N1 latency for each response site after stimulating all other electrode pairs (Fig. 2). We find that N1 latency correlates negatively with age in 14 out of 16 connections between temporal, central, parietal, and frontal regions (Spearman’s ρ, p_FDR corrected_<0.05). The number of CCEPs does not change with age indicating no change in overall level of connectivity, see Fig S2. The decrease in latency with age is observed across long-range connections (e.g. temporal - parietal), as well as short-range connections (e.g. temporal - temporal). This indicates that the human brain shows widespread decreases in transmission times with development.

**Fig. 2.**
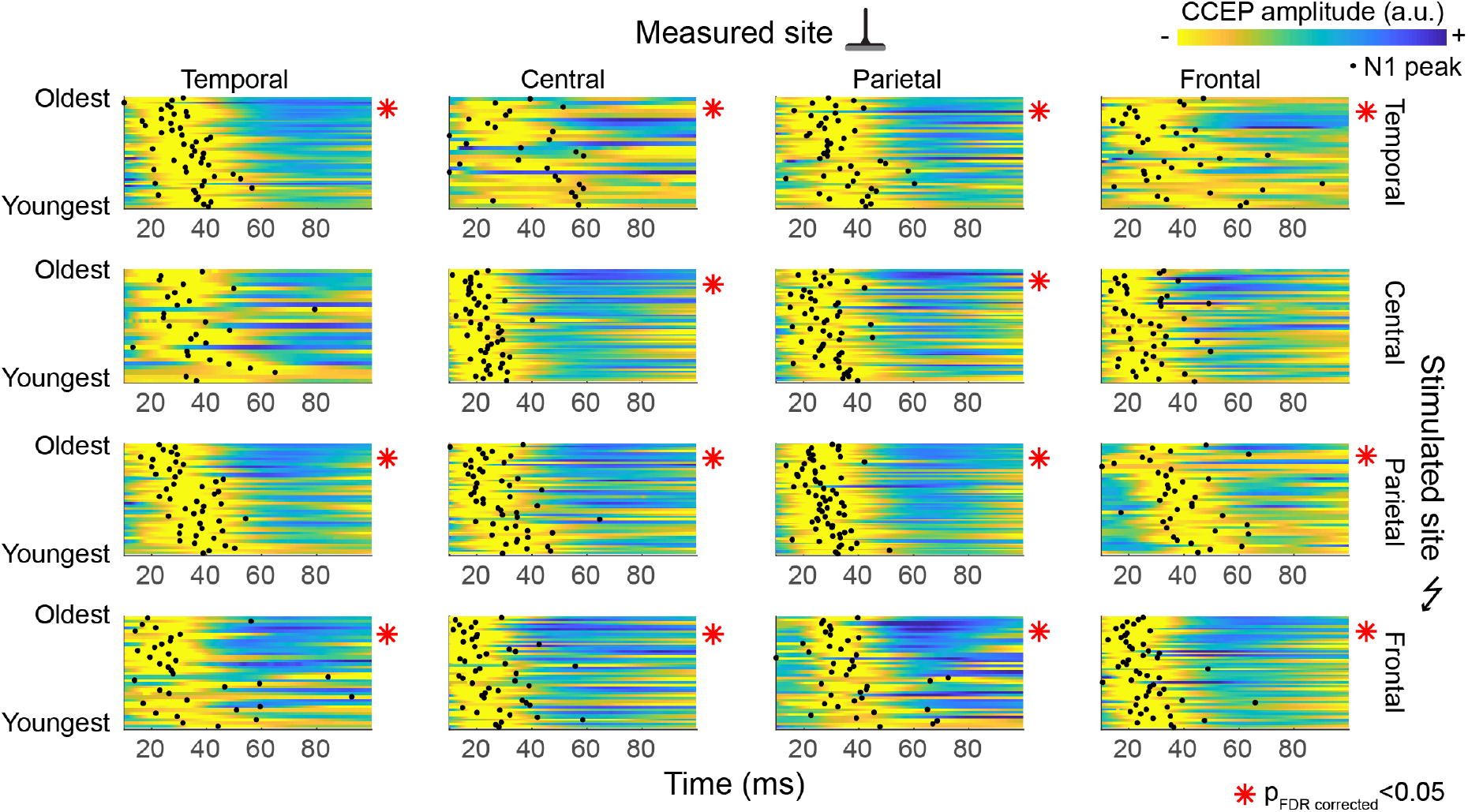
Normalized CCEP responses and N1 peak latency organized by age. Average CCEP responses per subject between temporal, central, parietal and frontal connections. CCEPs are unit length normalized and yellow indicates the largest negative deflection. A red * indicates a significant negative correlation between age and N1 latency (Spearman’s ρ, p_FDR corrected_<0.05). Black dots indicate the average N1 latency per subject.

We then estimate the latency change per year and fit a minimum to understand when this process reaches maturity across some of the well described long-range, structural connections. Frontal and parietal connections are largely governed by the superior longitudinal fasciculus; frontal and temporal areas are connected through the arcuate and uncinate fasciculi and temporal and parietal areas are connected by the posterior arcuate fasciculus (*19*). We fit a first and second order polynomial model where age predicts N1 latency (Fig. 3). These models have been used before in MRI studies of development (*11, 20*). Fitting these models with leave-one-out cross-validation further ensures that a single subject does not drive the results and lets the data indicate which connections are better described by a linear model or a quadratic model with a local minimum.

**Fig. 3.**
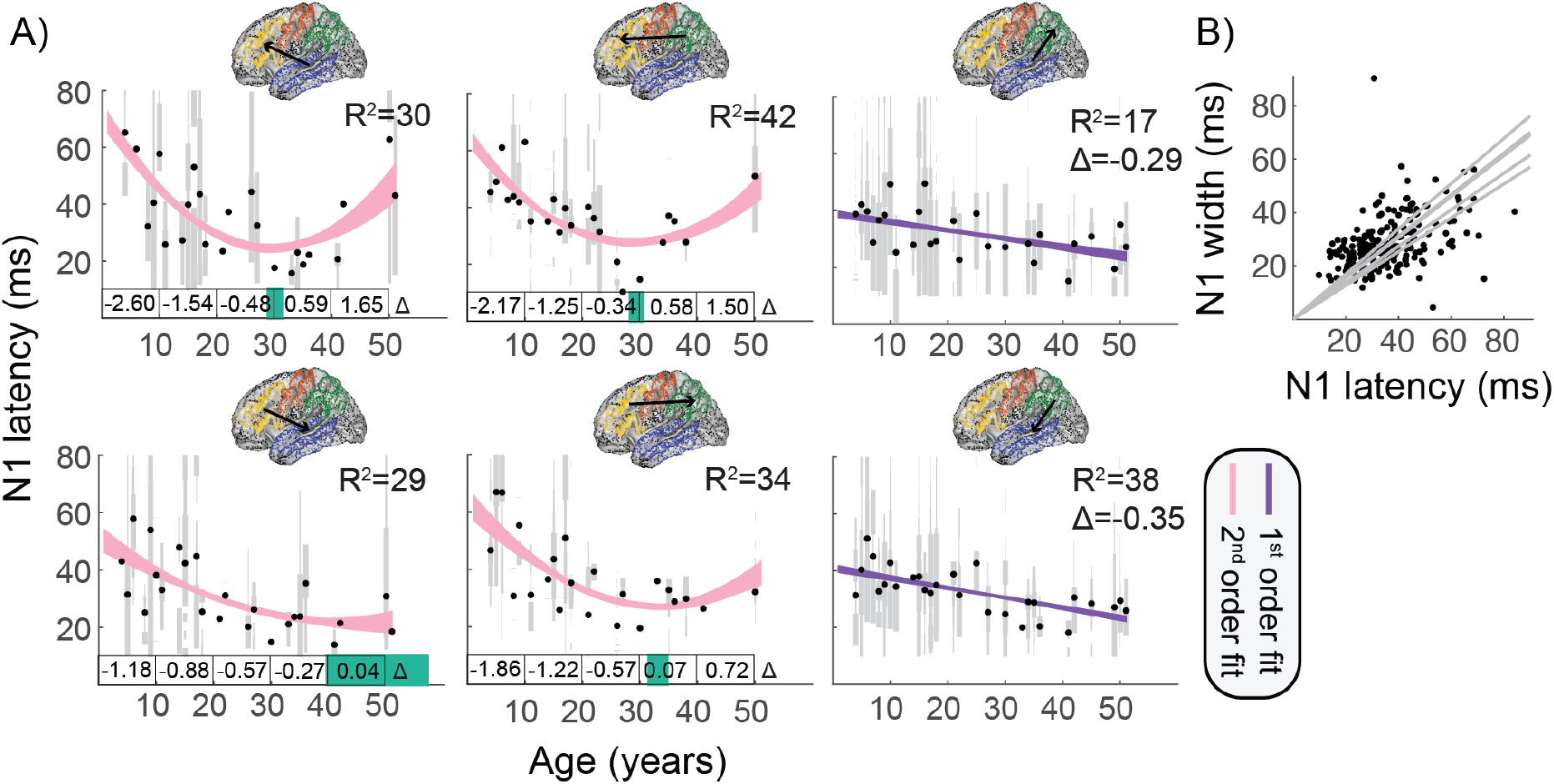
Long-range connections decrease in transmission delay with age. Average transmission delays estimated by the N1 latency between temporal, parietal and frontal electrodes. (**A)** Gray bars show distributions within each subject, bar width scales with the number of measured responses. Black dots show N1 latency averaged across subjects of the same age. First and second order polynomial models (shown with 95% confidence interval) explain the changes in N1 latency as a function of age. Explained variance (R^2^) calculated with leave-one-out cross-validation indicate whether the first order (purple) or second order (pink) polynomial model explains more variance. For second order polynomial model fits, the 95% confidence interval of the minimum is shown on the x-axis in green. For the first order polynomial fits, insets show the slope Δ in ms/year. For the second order polynomial fits, the slope Δ is displayed in ms/year averaged across ten years of age. (**B)** Across all 6 connections N1 latency correlates significantly with N1 width estimated by full width half max.

Across the long-range connections, age explains an average of 32% of variance in latency across subjects. A quadratic model best describes changes in transmission speed across frontal-parietal and frontal-temporal connections. Before age 10, latency decreases rapidly (by 1.2-2.6 ms/year), while between age 10 and 20 years, latency decreases at a slower rate (by 0.9-1.5 ms/year). A minimum average latency of 21–27 ms is reached around 34 years (range 29-44 years). A linear model better describes the maturation process of temporal-parietal connections, where latency decreases with ^~^0.32 ms/year. These small, yearly changes in conduction time result in a 1.5-2 fold increase in speed from childhood to middle age. This indicates that transmission speed across long-range connections matures well throughout adolescence.

Short-range connections across neighboring gyri such as the pre- and post-central gyrus and within frontal, temporal and parietal regions are supported by U-fibers. Figure 4 shows that latencies for these short connections, while already faster at an early age, decrease in a manner comparable to long-range connections. A quadratic model explains 43% of the variance in latency within both frontal and parietal regions. Up to age 10, latency within these two regions decreases rapidly (by ^~^0.85 ms/year), from age 10 to 20 latency decreased at a slower rate (by ^~^0.58 ms/year). A minimum latency (of ^~^23 ms) was reached between 35 and 38 years of age. Age related changes in temporal and central connections are better described by a linear model that predicts a yearly decrease of 0.26 ms. This linear model explains on average 21% of variance. This indicates that transmission speed across shortrange connections matures well throughout adolescence.

**Fig. 4.**
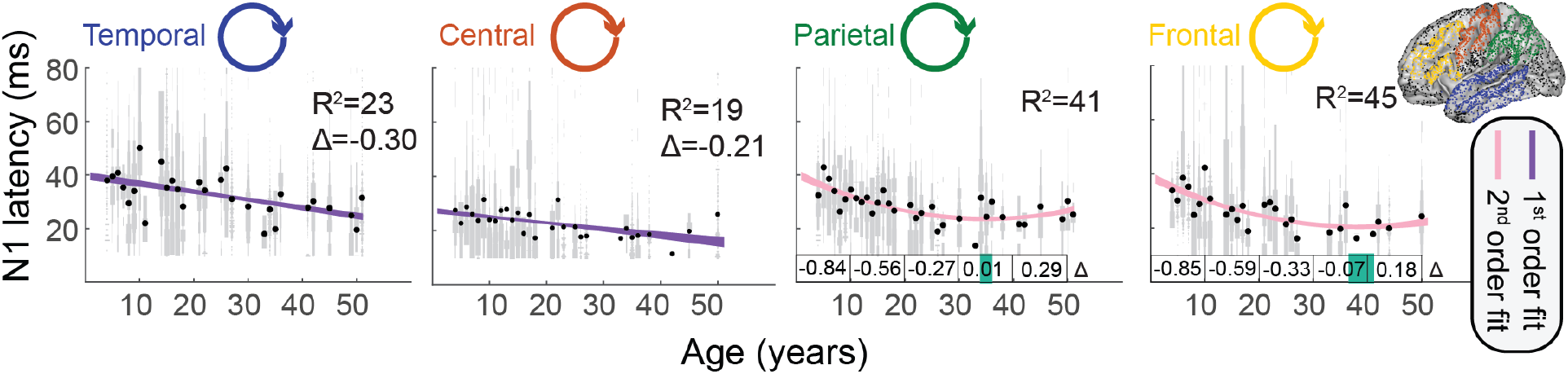
Short-range connections decrease in transmission delay with age. Average transmission delays estimated by the N1 latency within temporal, central, parietal and frontal electrodes. Gray bars show distributions within each subject, bar width scales with the number of measured responses. Black dots show N1 latency averaged across subjects of the same age. First and second order polynomial models (shown with 95% confidence interval) explain the changes in N1 latency as a function of age. Explained variance (R^2^) calculated with leave-one-out cross-validation indicate whether the first order (purple) or second order (pink) polynomial model explains more variance. For second order polynomial model fits, the 95% confidence interval of the minimum is shown on the x-axis in green. For the first order polynomial fits, insets show the slope Δ in ms/year. For the second order polynomial fits, the slope Δ is displayed in ms/year averaged across ten years of age.

Decreases in latency suggest faster timing of network interactions, but the precision of cortico-cortical communication also depends on variance between and within connections. We found no evidence for a relation between subject’s age and variance in latency between connected sites (p_FDR corrected_<0.05, see Fig. S3). However, we find that faster connections also had reduced variance between different connected regions sites (Fig. S4) and that faster N1 responses had smaller widths (Fig. 3B and Fig. S5). For many natural processes, the ratio between the mean and variance is fixed (*21*), and this principle seems to apply to the transmission speed of cortico-cortical connections with faster connections having more precise timing.

## Discussion

Our data indicate that transmission speeds are still maturing during adolescence and early adulthood. Many psychopathologies, like schizophrenia, anxiety disorders, depression, and bipolar disorders, can emerge during these periods (*13*), emphasizing the potential importance of our findings for these diseases. We note that, while our subjects suffered from epilepsy, there were no consistent effects of the seizure onset region on latency (Fig. S6, S7), and epilepsy may merely have added noise to the estimates. The large number of subjects this allows us to establish a normative baseline to which different pathologies may be compared.

A long maturation process of transmission speed aligns with findings from non-invasive neuroimaging studies that show that white matter pathways in the human brain mature well into early adulthood. The structure of the white matter pathways changes in terms of axon diameter as well as myelination (*22*). Conduction velocity in myelinated fibers increases with axon diameter, which ranges from 0.16 to 9 μm in the cortex (*23*). Quantitative MRI studies of the white matter pathways have captured some of these processes and show that white matter development follows a quadratic function with a peak between 30 and 40 years of age (*11*). This trajectory is comparable to the developmental trajectory of transmission speed that is shown in our data. These findings indicate that the long developmental trajectory of white matter axons can have a large effect on the timescales of neuronal communication in the human brain.

A simple characterization of the timing of direct cortico-cortical interactions has large implications for the temporal dynamics of brain function. Functional networks measured with resting state fMRI mature across the life-span (*24*), and graph theoretical analysis of these resting state fMRI networks have suggested that global efficiency increases from age 0 to 18 years (*25*). While the electrophysiological dynamics underlying these types of resting state fMRI changes remain unclear, decreases in latencies between long range connections may make it easier to transmit information. Neurons integrate information over time, as has been modeled by drift diffusion models for firing rates (*26*), as well as temporal receptive windows for fMRI and ECoG recordings (*27*). In drift diffusion models, a decision is made when activity reaches a certain threshold. With developing speed of connectivity within higher order areas, faster types of inputs may contribute relatively stronger to a decision at younger ages compared to adulthood. In temporal receptive windows, different brain regions are proposed to integrate information across various segments of time. Faster connectivity will allow more information to be integrated into a particular window, whereas slower connectivity will result in information being segregated.

Previous studies provide ways to estimate changes in velocity across the white matter axons with age. We calculated N1 peak latencies, but the earliest orthodromic responses likely arrive at the onset of the N1 response (*28*). Given the observed relation between latency and the width of the CCEPs, we can estimate onset decreases of the N1 responses from 25 to 12 ms from the N1 peak latency decreases from at 50 to 25 ms. If we assume a fiber length of 10 cm, an decreased latency of 25 to 12 ms would mean a velocity increase of 4 m/s to 8.4 m/s, which are typical conduction speeds for primate brain axons (*29*).

These two-fold increases in the speed of communication in the human connectome are observed in long-range as well as short-range connections. The large, consistent effects of age on transmission speed across sites provide normative estimates for the speed of cortico-cortical signaling across the life-span in distributed as well as local networks.

## Methods

### Subjects

All subjects who underwent epilepsy surgery in the UMC Utrecht between 2008 and 2020 have been included in a retrospective epilepsy surgery database (*30*), with approval of the Medical Ethical Committee from the UMC Utrecht. For subjects included from January 2008 and December 2017, the Medical Research Ethical Committee from the UMC Utrecht waived the need to ask for informed consent. Since January 2018, we explicitly ask subjects informed consent to collect their data for research purposes. When subjects underwent Single Pulse Electrical Stimulation (SPES) for clinical purposes during the intracranial grid monitoring period between 2012 and 2020, and electrode positions were determined from CT co-registered with T1 weighted MRI (*31*), we included these subjects in this study. In total, 75 subjects were included (median age: 17 years (4-51 years), 38 females).

During the evaluation for epilepsy, the seizure onset zone (SOZ) and eloquent cortex are delineated and a resection area is suggested to the surgeon. In 30 subjects, the seizure onset zone was added to the database by a clinical neurophysiologist to allow a comparison between latencies in and outside of the seizure onset region.

### Acquisition

Long-term ECoG data were recorded with subdural electrode grids and strips of 4.2 mm^2^ contact surface and an interelectrode distance of 1 cm (Ad-Tech, Racine, WI and PMT). Additional depth electrodes were implanted in 4 subjects but are not included in analyses.

SPES was performed during ECoG recordings with data sampled at 2048 Hz using a MicroMed LTM64/128 express EEG headbox with integrated programmable stimulator (MicroMed, Mogliano—Veneto, Italy). Ten monophasic stimuli with a pulse width of 1 ms were applied at a frequency of 0.2 Hz to two adjacent electrodes. A current intensity of 8 mA was used, but in case electrodes were located near central nerves or in the primary sensorimotor cortex, the intensity was lowered to 4 mA to avoid pain or twitches.

### Detecting N1-peak latency

For each electrode, 10 epochs with a time window of 2 s pre-stimulus to 3 s post-stimulus, time-locked to the stimulus, are corrected for baseline (median signal in a time window of 900 ms prior to stimulation (−1s to −0.1 s) and averaged for each stimulus pair (*32*). For each averaged epoch, the median is subtracted (−2s to −0.1 s), and the standard deviation (SD) is calculated in this pre-stimulus window. N1s are detected when the evoked response exceeds 3.4*SD in a time window of 9-100 ms post-stimulation, excluding earlier times due to potential stimulation artifacts. Electrodes which overlap with another grid, or are noisy, are excluded from detection.

### Electrode localization

We used Freesurfer to label the electrodes according to the cortical anatomy using the Destrieux atlas (*33*). We categorized electrodes based on location on the temporal lobe, frontal lobe, parietal lobe, occipital lobe, or central area, consisting of the pre- and post-central gyrus (Supplementary Table S1), as these regions are roughly connected through several large structural connections. For visualization purposes, we converted the individual subject’s electrode positions to MNI152 space. Regions without sufficient electrode coverage for across-subject correlations, such as the occipital lobe, were excluded from analyses.

## Data and code availability

The data that support the findings of this study will be made available in BIDS format on OpenNeuro. Code to analyse the data and generate all figures of this manuscript is available on GitHub: https://github.com/MultimodalNeuroimagingLab/mnl_ccepBids

## Acknowledgments

We thank the RESPect database group (K.P.J. Braun, C.H. Ferrier, T.A. Gebbink, P.H. Gosselaar, N.E.C. van Klink, M.A. van ‘t Klooster, J.M. Ophorst, P.C. van Rijen, E.V. Schaft, S.M.A. van der Salm, D. Sun, A. Velders and G.J.M. Zijlmans) for their contributions and help in collecting the data and Gabriella Ojeda Valencia for proof reading the manuscript. Research reported in this publication was supported by the National Institute of Mental Health of the National Institutes of Health under Award Number R01MH122258 (DH, FSSL, the content is solely the responsibility of the authors and does not necessarily represent the official views of the National Institutes of Health), the EpilepsieNL under Award Number NEF17-07 (DvB) and the UMC Utrecht Alexandre Suerman MD/PhD Stipendium 2015 (WZ).

## References

1. S. A. Neymotin, D. S. Daniels, B. Caldwell, R. A. McDougal, N. T. Carnevale, M. Jas, C. I. Moore, M. L. Hines, M. Hämäläinen, S. R. Jones, Human neocortical neurosolver (HNN), a new software tool for interpreting the cellular and network origin of human MEG/EEG data. Elife. 9 (2020), doi:10.7554/ELIFE.51214.

2. C. J. Honey, T. Thesen, T. H. Donner, L. J. Silber, C. E. Carlson, O. Devinsky, W. K. Doyle, N. Rubin, D. J. Heeger, U. Hasson, Slow cortical dynamics and the accumulation of information over long timescales. Neuron. 76, 423–434 (2012).

3. J. I. Gold, M. N. Shadlen, The neural basis of decision making. Annu. Rev. Neurosci. 30, 535–574 (2007).

4. R. Ratcliff, G. McKoon, Drift Diffusion Decision Model: Theory and data. Neural Comput. 20, 873–922 (2008).

5. G. Buzsáki, N. Logothetis, W. Singer, Scaling brain size, keeping timing: Evolutionary preservation of brain rhythms. Neuron. 80, 751–764 (2013).

6. Y. Mahajan, G. McArthur, Maturation of visual evoked potentials across adolescence. Brain Dev. 34, 655–666 (2012).

7. M. A. Crognale, Development, maturation, and aging of chromatic visual pathways: VEP results. J. Vis. 2, 438–450 (2002).

8. E. S. Orlrich, A. B. Barnet, I. P. Weiss, B. L. Shanks, Auditory evoked potential development in early childhood: A longitudinal study. Electroencephalogr. Clin. Neurophysiol. 44, 411–423 (1978).

9. J. Lemaréchal, M. Jedynak, L. Trebaul, A. Boyer, M. Bhattacharjee, P. Deman, V. Tuyisenge, L. Ayoubian, B. Chanteloup-forêt, C. Saubat, R. Zouglech, G. C. Reyes, S. Tourbier, P. Hagmann, C. Adam, C. Barba, T. Blauwblomme, J. Curot, F. Dubeau, M. Garcés, E. Hirsch, E. Landré, S. Liu, A brain atlas of axonal and synaptic delays based on modelling of cortico-cortical evoked potentials. Brain. awab362, 2–37 (2021).

10. B. Emery, Regulation of oligodendrocyte development. Science (80-.). 330, 779–782 (2010).

11. J. D. Yeatman, B. A. Wandell, A. A. Mezer, Lifespan maturation and degeneration of human brain white matter. Nat. Commun. 5, 4932 (2014).

12. T. Klingberg, C. J. Vaidya, J. D. E. Gabrieli, M. E. Moseley, M. Hedehus, Myelination and organization of the frontal white matter in children: A diffusion tensor MRI study. Neuroreport. 10, 2817–2821 (1999).

13. T. Paus, M. Keshavan, J. N. Giedd, Why do many psychiatric disorders emerge during adolescence? Nat Rev Neurosci. 9, 947–957 (2008).

14. R. Matsumoto, D. R. Nair, E. LaPresto, I. Najm, W. Bingaman, H. Shibasaki, H. O. Luders, H. O. Lüders, H. O. Luders, Functional connectivity in the human language system: A cortico-cortical evoked potential study. Brain. 127, 2316–2330 (2004).

15. U. Mitzdorf, Current source-density method and application in cat cerebral cortex: Investigation of evoked potentials and EEG phenomena. Physiol. Rev. 65, 37–100 (1985).

16. K. J. Miller, K.-R. Muller, D. Hermes, Basis profile curve identification to understand electrical stimulation effects in human brain networks. J. Clin. Transl. Sci. 5, 8–8 (2021).

17. C. J. Keller, S. Bickel, L. Entz, I. Ulbert, M. P. Milham, C. Kelly, A. D. Mehta, Intrinsic functional architecture predicts electrically evoked responses in the human brain. Proc. Natl. Acad. Sci. 108, 17234–17234 (2011).

18. R. S. Desikan, F. Ségonne, B. Fischl, B. T. Quinn, B. C. Dickerson, D. Blacker, R. L. Buckner, A. M. Dale, R. P. Maguire, B. T. Hyman, M. S. Albert, R. J. Killiany, An automated labeling system for subdividing the human cerebral cortex on MRI scans into gyral based regions of interest. Neuroimage. 31, 968–980 (2006).

19. D. Bullock, H. Takemura, C. F. Caiafa, L. Kitchell, B. McPherson, B. Caron, F. Pestilli, Associative white matter connecting the dorsal and ventral posterior human cortex. Brain Struct. Funct. 224, 2631–2660 (2019).

20. J. Blumenthal, J. Jeffries, E. X. Castellanos, H. Lin, A. Zidjdenbos, T. Paurs, A. C. Evans, J. L. Rapaport, J. N. Giedd, Brain development during childhood and adolescence: A longitudinal MRI study. Nat. Neurosci. 10, 861–863 (1999).

21. W. Gerstner, W. M. Kistler, R. Naud, L. Paninski, Neuronal Dynamics (Cambridge University Press, Cambridge, 2014).

22. T. Paus, Growth of white matter in the adolescent brain: Myelin or axon? Brain Cogn. 72, 26–35 (2010).

23. D. Liewald, R. Miller, N. Logothetis, H. J. Wagner, A. Schüz, Distribution of axon diameters in cortical white matter: an electron-microscopic study on three human brains and a macaque. Biol. Cybern. 108, 541–557 (2014).

24. N. U. F. Dosenbach, B. Nardos, A. L. Cohen, D. A. Fair, J. D. Power, J. A. Church, S. M. Nelson, G. S. Wig, A. C. Vogel, C. N. Lessov-Schlaggar, K. A. Barnes, J. W. Dubis, E. Feczko, R. S. Coalson, J. R. Pruett Jr., D. M. Barch, S. E. Petersen, B. L. Schlaggar, Prediction of Individual Brain Maturity Using fMRI. Science (80-.). 329, 1358–1361 (2010).

25. E. Gozdas, S. K. Holland, M. Altaye, C. A. Consortium, Developmental changes in functional brain networks from birth through adolescence. Hum. Brain Mapp., 1–11 (2018).

26. M. N. Shadlen, W. T. Newsome, Neural basis of a perceptual decision in the parietal cortex (area LIP) of the rhesus monkey. J. Neurophysiol. 86, 1916–1936 (2001).

27. C. J. Honey, T. Thesen, T. H. Donner, L. J. Silber, C. E. Carlson, O. Devinsky, W. K. Doyle, N. Rubin, D. J. Heeger, U. Hasson, Slow cortical dynamics and the accumulation of information over long timescales. Neuron. 76, 423–434 (2012).

28. R. Matsumoto, T. Kunieda, D. Nair, Single pulse electrical stimulation to probe functional and pathological connectivity in epilepsy. Seizure. 44, 27–36 (2017).

29. G. M. Innocenti, A. Vercelli, R. Caminiti, The diameter of cortical axons depends both on the area of origin and target. Cereb. Cortex. 24, 2178–2188 (2014).

30. M. Demuru, D. van Blooijs, W. Zweiphenning, D. Hermes, F. Leijten, M. Zijlmans, D. van Blooijs, K. P. J. Braun, M. Demuru, P. van Eijsden, C. H. Ferrier, T. A. Gebbink, P. H. Gosselaar, G. J. M. Huiskamp, N. E. C. van Klink, M. A. van t Klooster, F. S. S. Leijten, J. M. Ophorst, P. C. van Rijen, E. V. Schaft, S. M. A. van der Salm, D. Sun, A. Velders, G. J. M. Zijlmans, W. J. E. M. Zweiphenning, A practical workflow for organizing clinical intraoperative and long-term iEEG data in BIDS. medRxiv (2020), doi:10.1101/2020.12.07.20245290.

31. D. Hermes, K. J. Miller, H. J. Noordmans, M. J. Vansteensel, N. F. Ramsey, Automated electrocorticographic electrode localization on individually rendered brain surfaces. J. Neurosci. Methods. 185, 293–298 (2010).

32. D. van Blooijs, F. S. S. Leijten, P. C. van Rijen, H. G. E. Meijer, G. J. M. Huiskamp, D. van Blooijs, F. S. S. Leijten, P. C. van Rijen, H. G. E. Meijer, G. J. M. Huiskamp, Evoked directional network characteristics of epileptogenic tissue derived from single pulse electrical stimulation. Hum. Brain Mapp., 1–12 (2018).

33. C. Destrieux, B. Fischl, A. Dale, E. Halgren, Automatic parcellation of human cortical gyri and sulci using standard anatomical nomenclature. Neuroimage. 53, 1–15 (2010).

