## Supplementary Table S1 for "The development of efficient communication in the human connectome"

**This PDF file includes:**

Figs. S1 to S7
Table S1

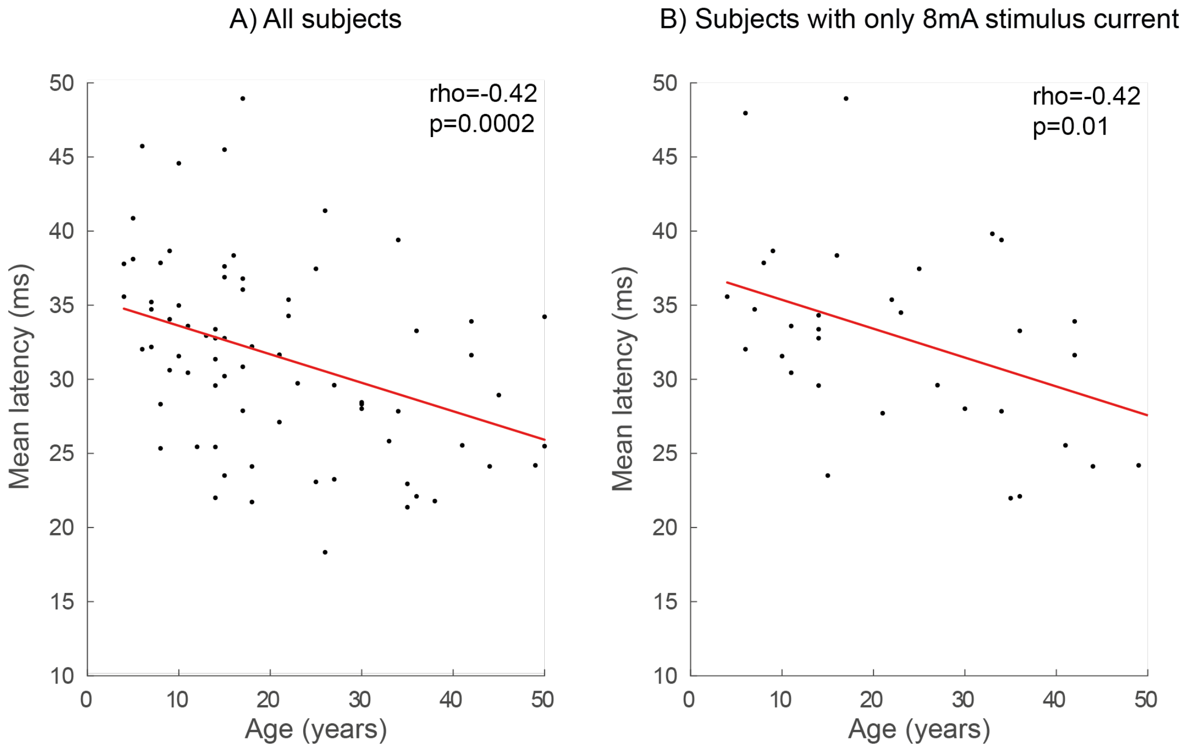


**Fig. S1. Similar relation between age (years) and latency (ms) at different stimulation currents.** To ensure that the relation between age and latency was not driven by the fact that some electrode pairs were stimulated with a current of 8 mA, while others had a current of 4 mA, we calculated the average latency per subject across all connected electrodes. **A)** The relation between age and latency across all subjects stimulated with 4 mA or 8 mA shows a negative relation between age and latency. **B)** The relation between age and latency across subjects where a stimulus current of 8 mA was applied shows a similar negative relation between age and latency.

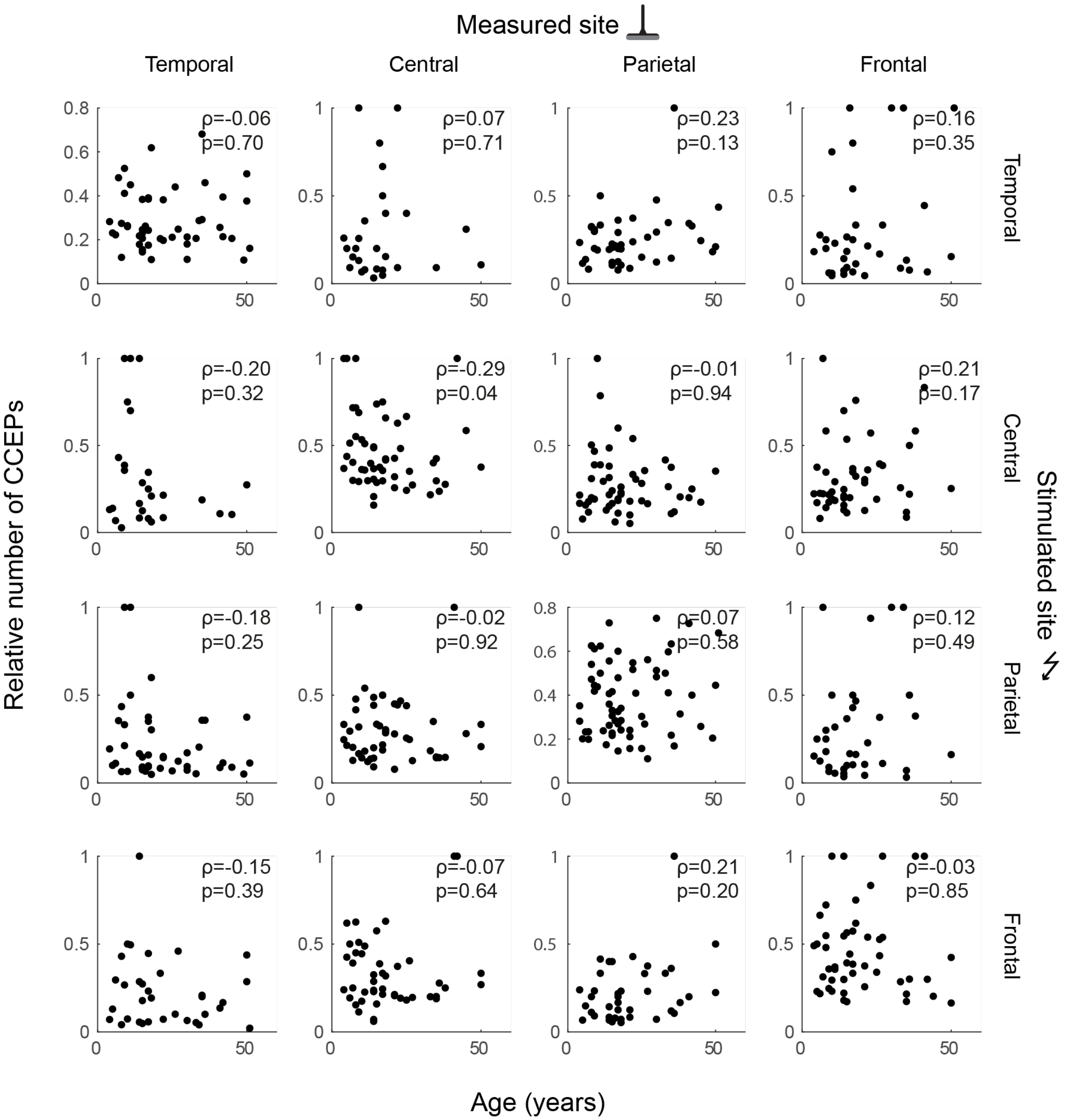


**Fig. S2. Relation between age and relative number of CCEPs.** Age (x-axis) versus the relative number of CCEPs in each region (y-axis). Since the number of electrodes on regions differ between subjects, we then divide the number of significant CCEPs by the total number of electrodes covering this region to calculate a relative number of CCEPs. We do not observe significant relations between age and the number of CCEPs (Spearman's ρ, $p_{FDR\ corrected}$<0.05).

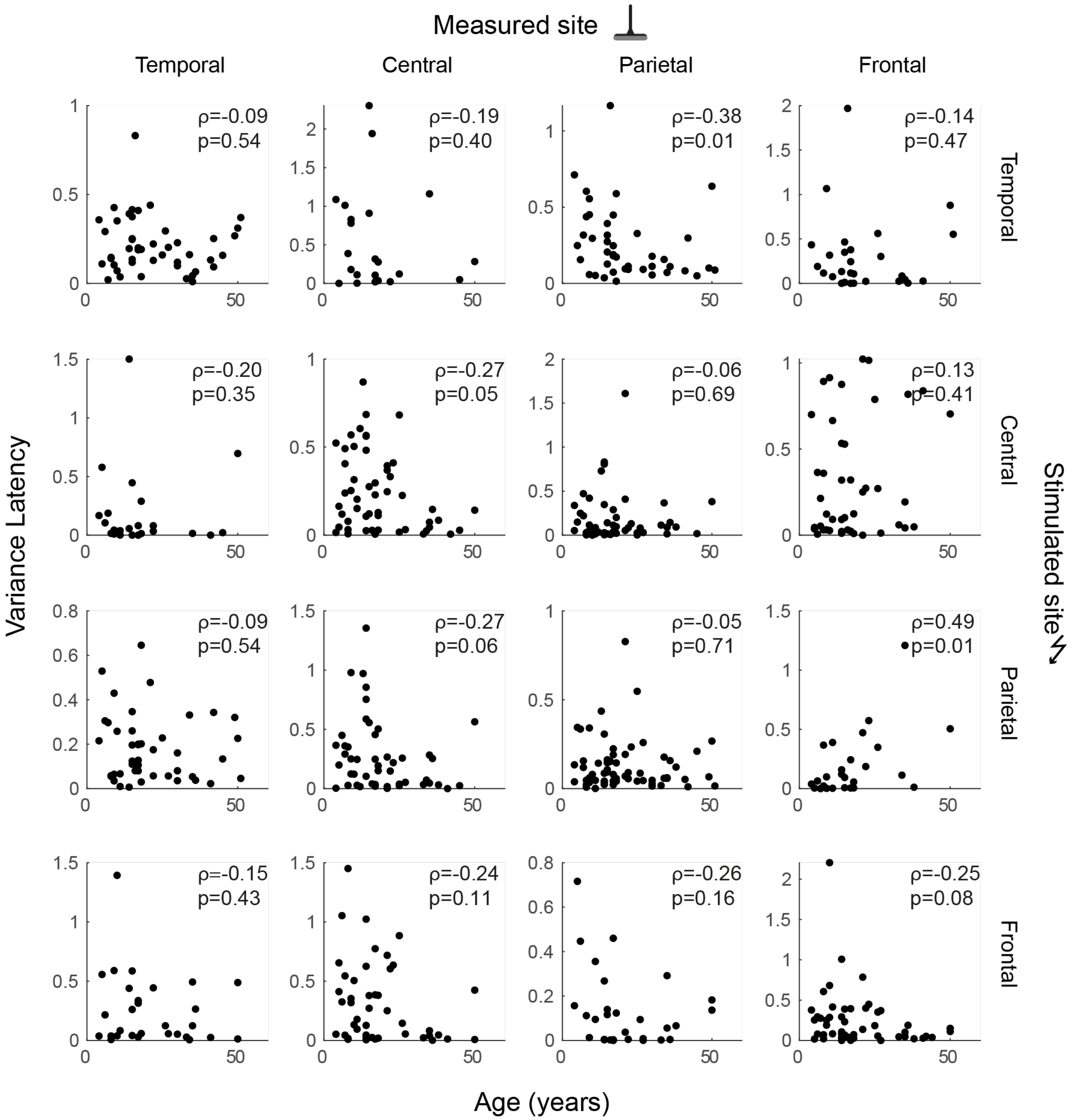


**Fig. S3. Relation between age (years) and variance in N1-latency across sites.** We do not observe significant correlations between age and variance in latency (Spearman's ρ, $p_{FDR\ corrected}<0.05$).

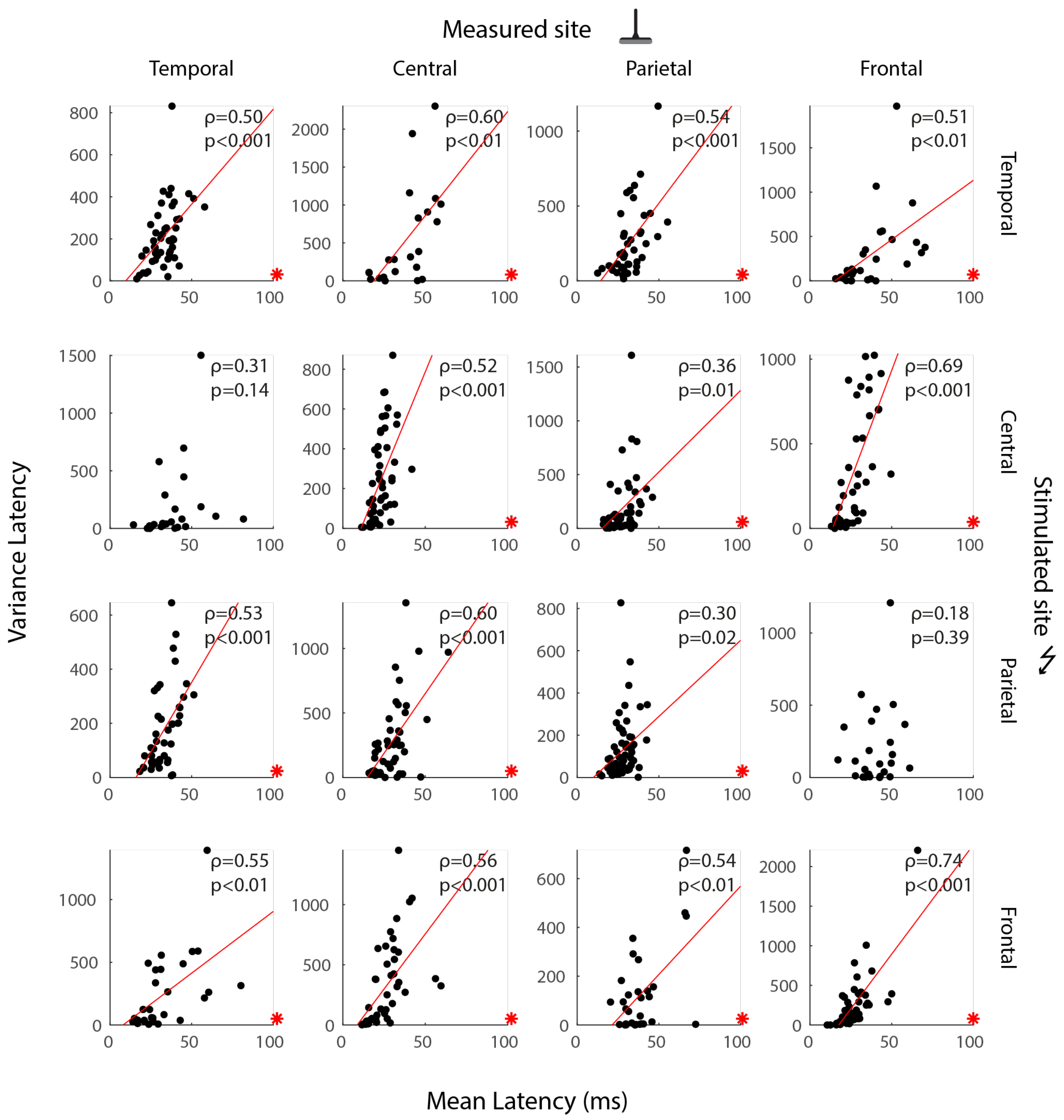


**Fig. S4. The relation between mean N1-latency (ms) and variance in latency across sites.** In 14 out of 16 connections we observe a positive relation between mean and variance in latency (Pearson's ρ, $p_{\text{FDR corrected}}<0.05$), red asterisk indicates significance. This shows that with increasing latency and faster connections, the timing across the different connected sites gets more similar.

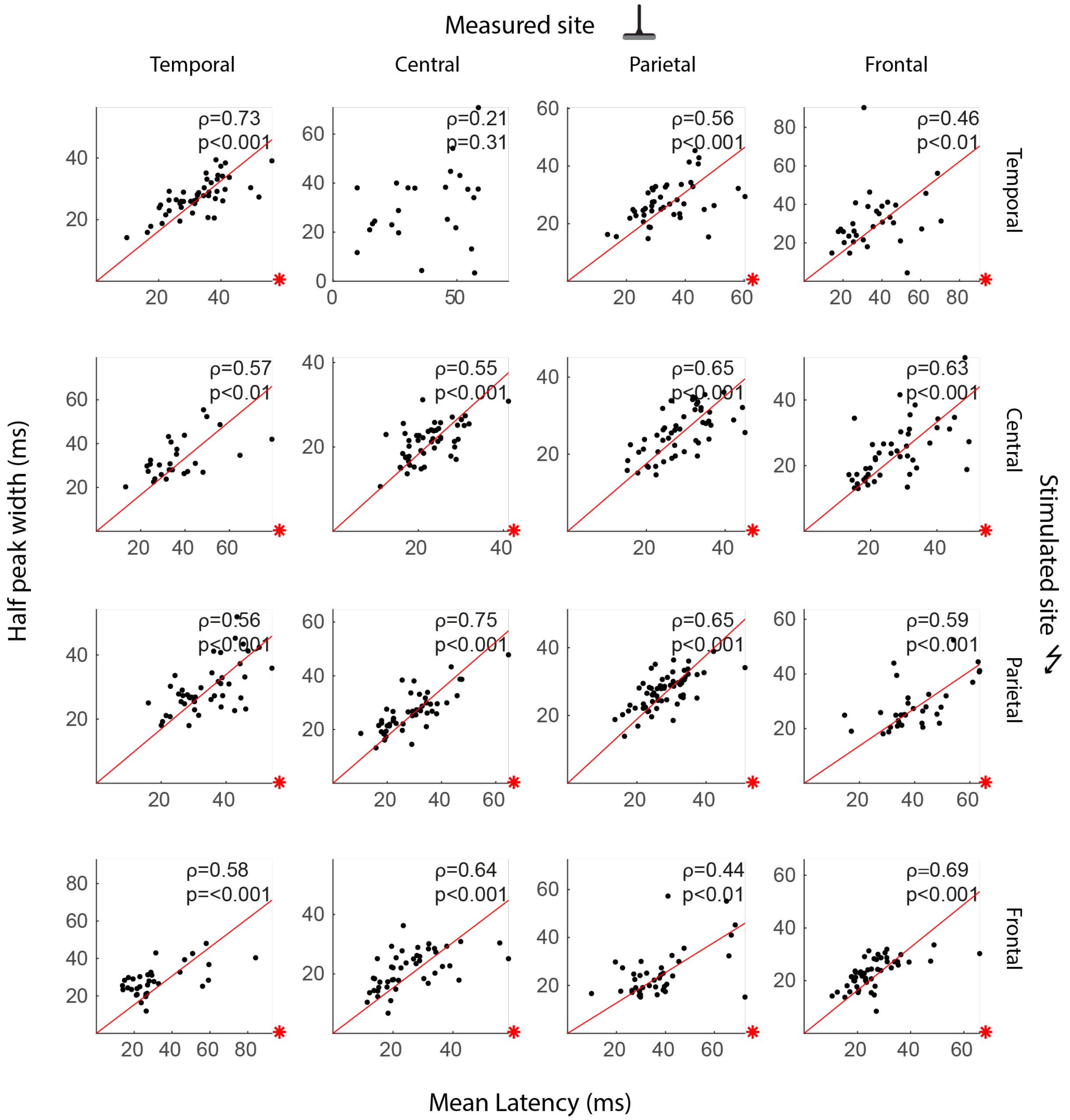


**Fig. S5. Relation between mean N1 latency (ms) and half N1 peak width (ms).** The variance in timing of one evoked potential could be calculated as the full width half max of the N1 peak, where the amplitude is 50% of the N1-peak amplitude. This could be considered a measure for the variance of synchronous signals arriving in the responding electrode. In 15 out of 16 connections, we observe a positive relation between the latency and the half N1 peak width (Spearman's ρ, $p_{\text{FDR corrected}}$<0.05), red asterisk indicates significance.

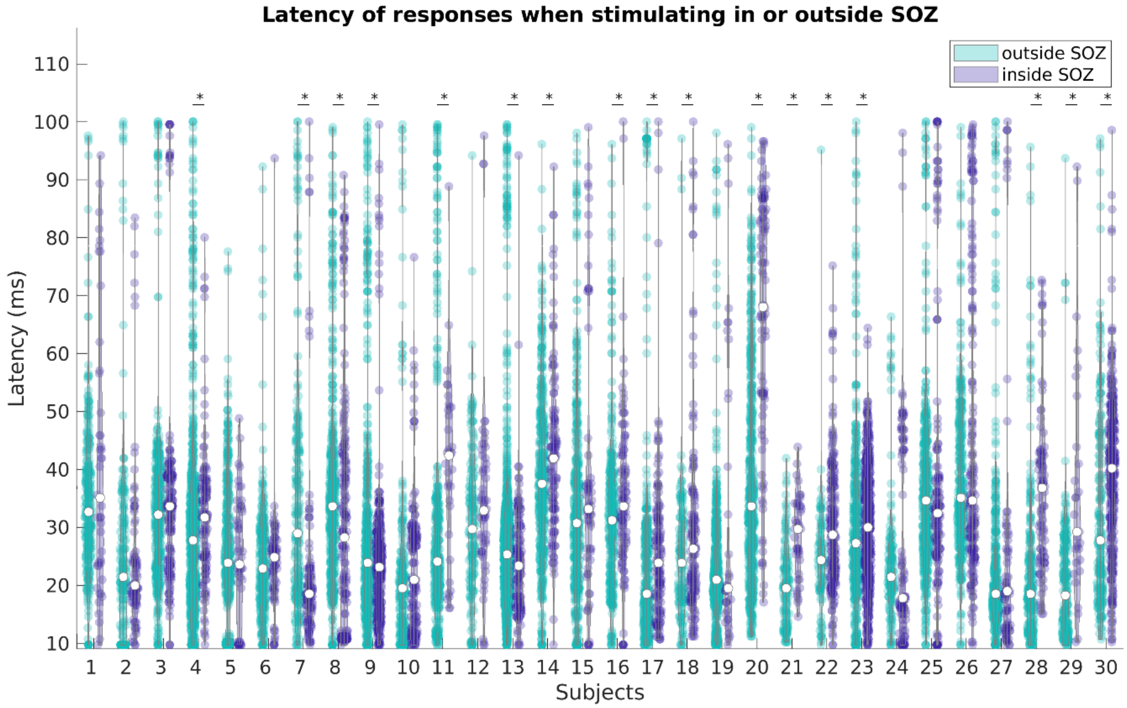


**Fig. S6. The latency of N1 peaks in response electrodes after stimulating in or outside the seizure onset zone (SOZ).** For 30 subjects, latencies for CCEPs measured after stimulating in or outside of the SOZ. The asterisks display the subjects in whom a significant difference is found after stimulating in or outside the SOZ (Mann Whitney U test, $p_{FDR\ corrected}<0.05$). In 13 subjects, we find an increase in latency in response electrodes when the SOZ is stimulated. In 4 subjects, we find a decrease in latency in response electrodes when the SOZ is stimulated.

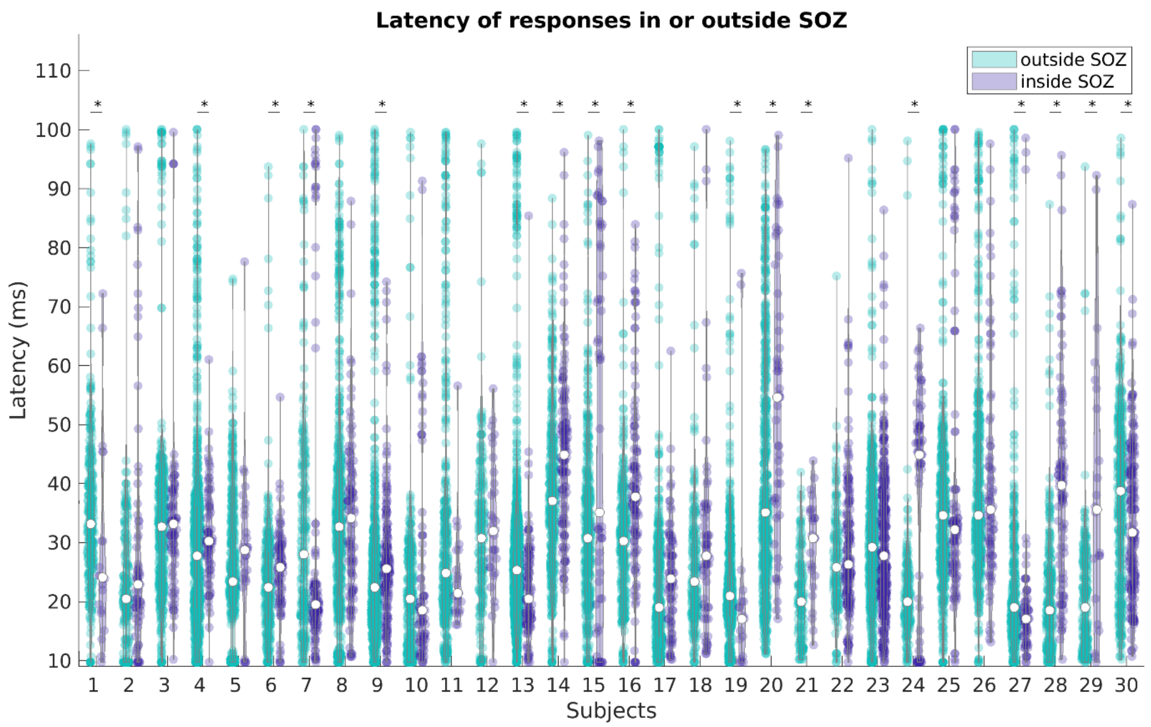


**Fig. S7. The latency of responses in or outside the seizure onset zone (SOZ) after stimulating elsewhere.** For 30 subjects, latencies for CCEPs in or outside of the SOZ when stimulating at other sites. The asterisks display the subjects in whom a significant difference is found in latencies in and outside SOZ (Mann Whitney U test, $p_{FDR\ corrected}<0.05$). In 6 subjects, the latency is decreased in the SOZ. In 11 subjects, the latency is increased in the SOZ.

| **Brain region** | **Destrieux regions** | **Total #electrodes in each region (median (min-max))** |
|---|---|---|
| Temporal | Temporal inferior gyrus, temporal middle gyrus, temporal superior lateral gyrus, occipito-medial-parahippocampal gyrus, occipito-temporal-lateral-fusiform gyrus | 1204 (10 (0-55)) |
| Frontal | Frontal inferior-triangular gyrus, frontal middle gyrus, frontal inferior-opercular gyrus | 771 (7(0-36)) |
| Parietal | Parietal inferior-angular gyrus, parietal inferior-supramarginalis gyrus, parietal superior gyrus | 891 (11 (0-34)) |
| Occipital | Occipital middle gyrus, occipital medial-lingual gyrus, occipital pole | 269 (0 (9-23)) |
| Central | Precentral gyrus, post central gyrus, central sulcus | 691 (7 (0-31)) |

**Table S1. Overview of number of electrode positions in different brain regions.** Electrodes were assigned to several brain regions based on the label from the Destrieux atlas in Freesurfer.
